# Unraveling the start element and regulatory divergence of core promoters across the domain Bacteria

**DOI:** 10.1101/2025.01.23.634641

**Authors:** Syue-Ting Kuo, Joshua Kevin Chang, Clara Chang, Wei-Yi Shen, Christine Hsu, Sheng-Wen Lai, Hsin-Hung David Chou

**Author notes:** These authors contributed equally to this work (Syue-Ting Kuo, Joshua Kevin Chang, Clara Chang). Present address: Department of Biomedical Engineering, University of California Davis; Davis, 95616, USA. Present address: Department of Molecular and Cell Biology, University of California San Diego; San Diego, 92122, USA.

## Abstract

Core promoters comprise multiple elements whose interaction with RNA polymerase initiates transcription. Despite decades of research, substantial sequence and length variation of promoter elements has hindered efforts to elucidate their function and the evolutionary diversity of transcriptional regulation. Combining massively parallel assays, biophysical modeling, and functional validation, we systematically dissected the promoter architecture upstream of experimentally determined transcription start sites in 49 phylogenetically diverse bacterial genomes (GC content: 27.8–72.1%). We identified a conserved 3-bp promoter element, termed ‘start,’ that dictates transcription start site selection and enhances transcription. We uncovered a four-region organization within the variable spacer element, whose sequence composition modulates transcription by up to 600-fold. We showed that the discriminator element is conserved in Terrabacteria but diverse in Gracilicutes, the two major bacterial clades. High discriminator sequence diversity in Gracilicutes likely reflects diversifying evolution, enabling promoter-encoded regulation to orchestrate global gene expression in response to growth rate changes. Together, our findings reveal broad conservation of bacterial promoter organization while highlighting regulatory divergence of promoter elements and RNA polymerase between Terrabacteria and Gracilicutes. Sequence and functional similarities between bacterial promoter elements and their archaeal and eukaryotic counterparts further suggest a shared evolutionary origin of promoter architecture.

## Introduction

Genomes use just four nucleobases to encode myriads of genes and regulatory sequences. Deciphering this language of life is crucial — not only for understanding how organisms survive today but also for tracing the evolutionary history on Earth and projecting their future trajectories. Identifying genes’ coding sequences is straightforward because they consist of codons, adhere to the reading frame, and show evident homology across species. In contrast, regulatory sequences, particularly core promoters, comprise multiple elements varying in length and sequence composition, making them difficult to recognize, especially in non-model organisms^1^.

In bacteria, core promoters contain UP,-35, spacer, extended-10 (Ex),-10, and discriminator (Dis) elements in the 5′-to-3′ order (Fig. 1a)^2^. Transcription begins with RNA polymerase (RNAP) binding to the-35 and-10 elements via the interchangeable σ subunit, followed by unwinding of DNA at the-10 element to synthesize RNA from the downstream transcription start site (TSS)^3^. The-35 and-10 elements are both hexamers showing consensus sequences TTGACA and TATAAT, respectively. The spacer element, ranging from 15 to 19 bp, exhibits variable sequence composition and limited RNAP contact, with 17 bp as the preferred length for transcription^4–7^. The Ex element refers to a TG motif (or TRTG, commonly seen in Firmicutes; R = purine), appearing 2 bp upstream of the-10 element within the spacer element^8,9^. The UP element, bound by the C-terminal domain of the RNAP α subunit, displays alternating A-and T-tracts and lacks a defined length^10,11^. The Dis element, lying downstream of the-10 element, interacts with RNAP σ, β, and β′ subunits and shows high sequence variation^12–15^.

**Fig. 1.**
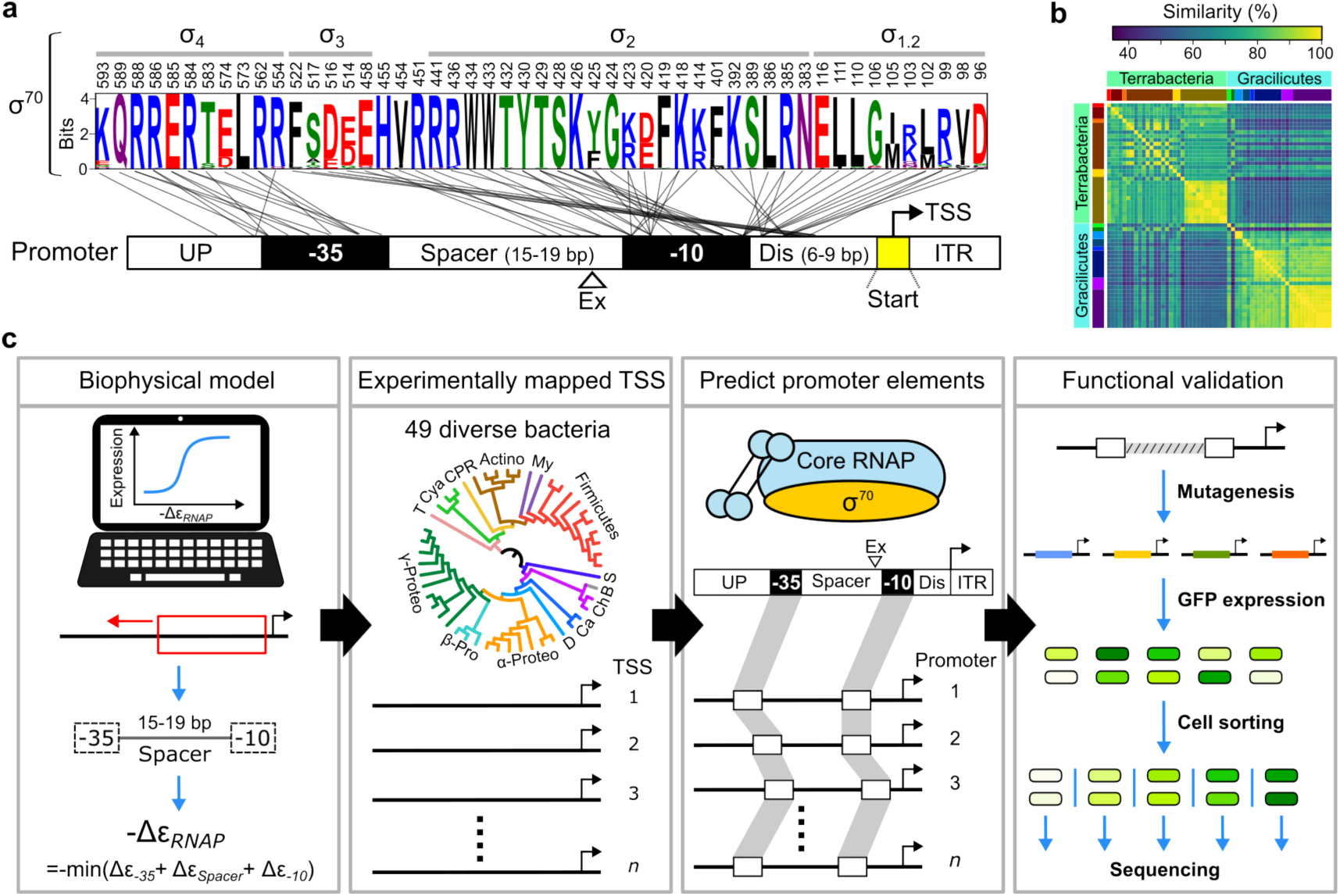
Dissecting the function, diversity, and evolution of bacterial promoter elements. **a**, σ^70^–promoter interaction. Promoter architecture highlights the broadly conserved start element, named in this study. Only σ^70^ residues that physically contact the promoter are shown^13,45,61,62^. Residue positions and their domain associations correspond to *E. coli* σ^70^. Sequence conservation among the 49 examined bacterial species is quantified by information content (sequence alignment shown in Extended Data Fig. 1a). ITR, initial transcribed region. **b**, Pairwise protein sequence similarity of the σ^70^ subunit among 49 species representing the two major bacterial clades. Species are ordered and lineages are distinguished by colors, as in Fig. 4a. Protein information and the maximum likelihood tree are shown in Extended Data Fig. 1. **c**, Research workflow, including model construction, collection of experimentally determined TSS, promoter element prediction, and functional validation.

Promoter evolution is shaped by multiple factors, including the corresponding σ subunit, genomic GC content, selection for gene expression, interaction with transcription factors, and functional complementation among promoter elements^2,16,17^. Decoding the regulatory syntax across diverse bacterial species is fundamental to advancing our understanding of transcriptional regulation and its evolution. Such knowledge forms the basis for developing predictive models that identify native promoters in genomes and guide the design of regulatory elements for synthetic biology applications. Earlier studies applied probabilistic models, like position weight matrices, and machine learning algorithms to reveal sequence features within individual promoter elements and across entire promoter regions^4,9,18–20^. More recently, researchers have applied massively parallel assays to explore the promoter sequence–function landscape by synthesizing and characterizing tens of thousands of promoter variants^21–25^. These efforts have enabled accurate identification of native promoters in model species (e.g., *Escherichia coli* and *Bacillus subtilis*) and yielded models that predict promoter strength in *E. coli* with impressive precision. To shed light on the diversity of promoters and gene regulation, future studies must look beyond model organisms^26–31^. However, progress has been hindered by limited empirical data, a wide range of genomic GC content, and substantial sequence and length variability among bacterial promoters—despite their being more compact and architecturally better defined than eukaryotic promoters^32^. Challenges in recognizing diverse bacterial promoters and their constituent elements have prompted researchers to reconsider the adequacy of the current promoter concept^16,2^.

Changes in gene expression are critical to adaptive evolution^33^, but limited knowledge of regulatory syntax has impeded our endeavor to unravel the underlying mechanisms. We reason that the field of gene regulation can be advanced through precise segmentation and alignment of promoter elements for experimental characterization and model development. Since the-35 and - 10 elements are relatively conserved and separate variable elements in bacterial promoters, we propose building an accurate-35 and-10 prediction model as the solution. We consider this approach feasible because residues on the primary σ subunit (σ^70^) that contact the-35 and-10 elements are highly conserved, even though the overall σ^70^ subunit has diverged significantly (Fig. 1a,b and Extended Data Fig. 1). To do so, we built a biophysical model trained on comprehensive sequence–function mapping of the-35 and-10 elements in *E. coli* (Fig. 2). The model was then applied to dissect core promoter organization upstream of experimentally determined TSS in 49 diverse bacterial genomes spanning the domain Bacteria (genome GC%: 27.8-72.1; Figs.1c, 3, 4a and Table 1). Subsequently, we performed experiments and analyses to validate the functional significance of the observed clade-specific and domain-wide promoter sequence conservation.

**Fig. 2.**
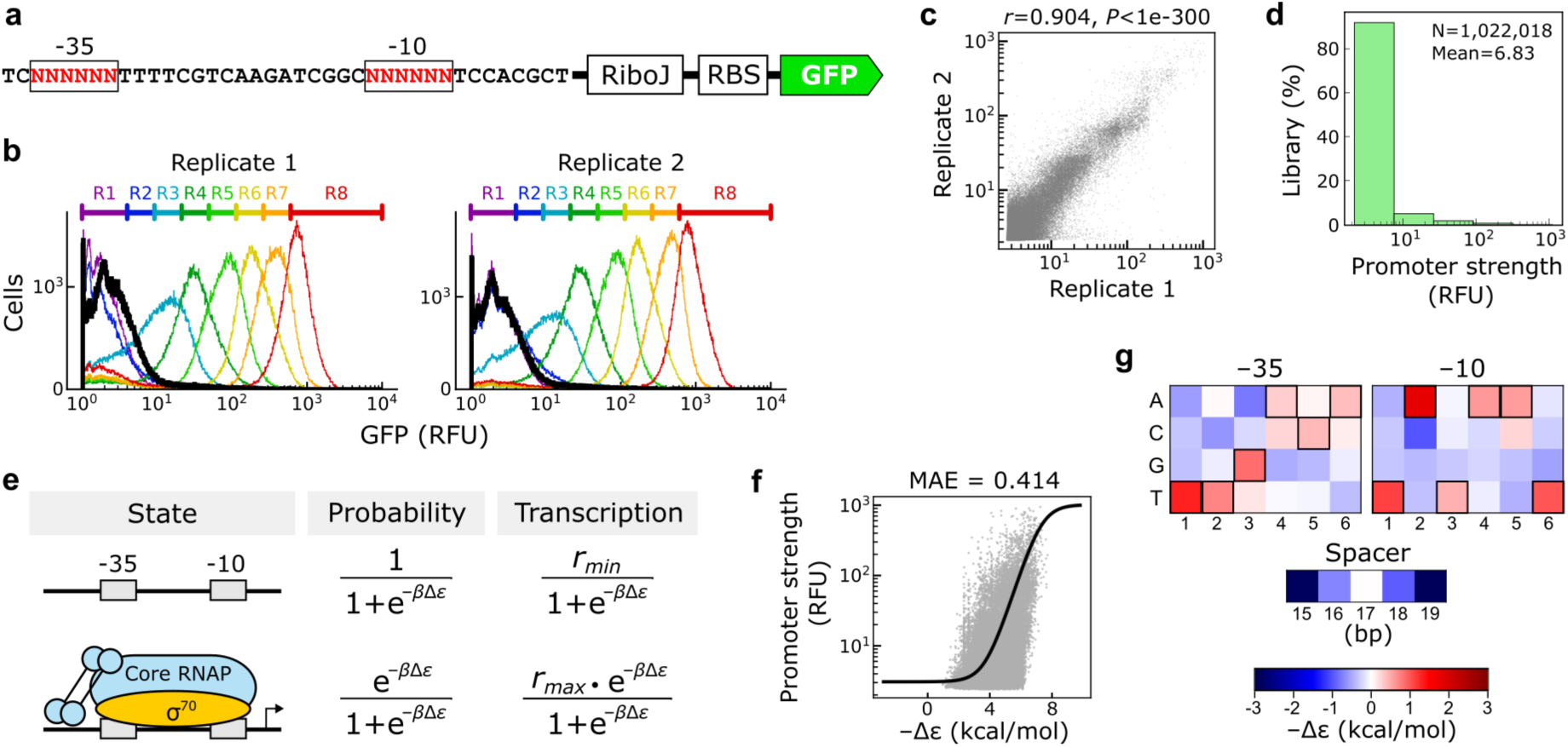
Determining the-35 and-10 sequence–function relationship for model construction. **a**, Promoter-reporter construct of the PL library. Two 6-bp randomized regions are marked in red. RiboJ^63^, self-cleaving ribozyme to make mRNA with a uniform 5′ end; RBS, ribosome binding site; GFP, green fluorescent protein. **b**, Quality of fluorescence-activated cell sorting in two sort-seq replicate experiments. The fluorescence distributions of eight sorting ranks (R1-R8) are distinguished by colors. The black curve indicates the whole-library distribution. RFU, relative fluorescence unit. **c**, Correlation (Pearson’s *r*, *t*-test) between promoter strength measured in two sort-seq replicates. **d**, Library distribution of promoter strength. **e**, Biophysical model of transcription. *−Δε_RNAP_*, RANP–promoter binding energy; *r_max_*, transcription rate constant; *r_min_*, basal measurement noise; *β = 1/RT*. **f**, Model fitting performance. The curve and dots represent model predictions and experimental data, respectively. MAE, mean absolute error. **g**, Model-estimated free energy (*−Δε*) contributions of each nucleobase in the-35 and-10 elements and of each spacer length. Bases representing the consensus sequences are outlined for emphasis.

**Table 1.** Bacterial genomes and TSS.

| Species | Acronym | Lineage | Genome accession no. | Experimentally mapped TSS |  |  |
| --- | --- | --- | --- | --- | --- | --- |
|  |  |  |  | Technology | Number | PMID |
| *Thermotoga maritima* | Th.ma | Thermotogota | CP004077.1 | dRNA-seq | 550 | 23637642 |
| *Anabaena* sp. | An.sp | Cyanobacteria | NC_003272.1 | dRNA-seq | 3955 | 22135468 |
| *Synechococcus elongatus* | Sy.el | Cyanobacteria | CP006471.1 | dRNA-seq | 2429 | 30127850 |
| *Synechocystis* sp. | Sy.sp | Cyanobacteria | NC_000911.1 | dRNA-seq | 170 | 21245330 |
| *Nanosynbacter lyticus* | Na.ly | CPR† | NZ_CP007496.1 | RNA-seq* | 188 | 35975994 |
| *Streptomyces coelicolor* | St.co | Actinobacteriota | NC_003888.3 | dRNA-seq | 1016 | 27251447 |
| *Corynebacterium diphtheriae* | Co.di | Actinobacteriota | NC_002935.2 | dRNA-seq | 1202 | 29370758 |
| *Corynebacterium glutamicum* | Co.gl | Actinobacteriota | BX927147.1 | dRNA-seq | 2147 | 24341750 |
| *Mycolicibacterium smegmatis* | My.sm | Actinobacteriota | NC_008596.1 | dRNA-seq | 3043 | 30984135 |
| *Mycobacterium tuberculosis* | My.tu | Actinobacteriota | AL123456.3 | dRNA-seq | 1778 | 24268774 |
| *Phytoplasma* sp. | Ph.sp | Mycoplasmata | AP006628.2 | dRNA-seq | 82 | 29039971 |
| *Mycoplasma pneumoniae* | My.pn | Mycoplasmata | NC_000912.1 | RNA-seq* | 718 | 22617959 |
| *Clostridioides difficile* | Cl.di | Firmicutes | NC_009089.1 | dRNA-seq | 1288 | 32903654 |
| *Lachnoclostridium phytofermentans* | La.ph | Firmicutes | NC_010001.1 | Capp-Switch | 2057 | 27982035 |
| *Paenibacillus riograndensis* | Pa.ri | Firmicutes | NZ_LN831776.1 | dRNA-seq | 1269 | 29100491 |
| *Bacillus subtilis* | Ba.su | Firmicutes | NC_000964.3 | dRNA-seq | 600 | 20525796 |
| *Staphylococcus epidermidis* | St.ep | Firmicutes | NC_004461.1 | dRNA-seq | 1206 | 27806702 |
| *Staphylococcus aureus* | St.au | Firmicutes | NC_003923.1 | dRNA-seq | 1800 | 27806702 |
| *Listeria innocua* | Li.in | Firmicutes | NC_003212.1 | dRNA-seq | 911 | 22617957 |
| *Listeria monocytogenes* | Li.mo | Firmicutes | NC_003210.1 | dRNA-seq | 1079 | 22617957 |
| *Streptococcus pyogenes* | St.py | Firmicutes | LR031521.1 | dRNA-seq | 683 | 30902048 |
| *Streptococcus agalactiae* | St.ag | Firmicutes | NC_004368.1 | dRNA-seq | 885 | 26024923 |
| *Leptospira interrogans* | Le.in | Spirochaetota | NZ_LT962963.1 | dRNA-seq | 2864 | 28154810 |
| *Bacteroides thetaiotaomicron* | Ba.th | Bacteroidota | NC_004663.1 | dRNA-seq | 1616 | 32678091 |
| *Chlamydia pneumoniae* | Ch.pn | Chlamydiota | NC_000922.1 | dRNA-seq | 382 | 21989159 |
| *Chlamydia trachomatis* | Ch.tr | Chlamydiota | NC_010280.1 | dRNA-seq | 349 | 19923228 |
| *Campylobacter jejuni* | Ca.je | Campylobacterota | NC_002163.1 | dRNA-seq | 675 | 23696746 |
| *Helicobacter pylori* | He.py | Campylobacterota | NC_000915.1 | dRNA-seq | 716 | 26091613 |
| *Geobacter sulfurreducens* | Ge.su | Desulfobacterota | NC_002939.5 | dRNA-seq | 875 | 27488344 |
| *Zymomonas mobilis* | Zy.mo | Alphaproteobacteria | NZ_CP023715.1 | dRNA-seq | 3205 | 32694125 |
| *Novosphingobium aromaticivorans* | No.ar | Alphaproteobacteria | NC_007794.1 | dRNA-seq | 944 | 34254822 |
| *Rhodobacter sphaeroides* | Rh.sp | Alphaproteobacteria | NC_007493.2 | dRNA-seq | 1087 | 34254822 |
| *Caulobacter crescentis* | Ca.cr | Alphaproteobacteria | NC_011916.1 | dRNA-seq | 1443 | 25569173 |
| *Agrobacterium tumefaciens* | Ag.tu | Alphaproteobacteria | NC_003062.2 | dRNA-seq | 468 | 23950988 |
| *Methylobacterium extorquens* | Me.ex | Alphaproteobacteria | NC_012988.1 | dRNA-seq | 2030 | 35889020 |
| *Bradyrhizobium japonicum* | Br.ja | Alphaproteobacteria | NC_004463.1 | dRNA-seq | 4451 | 27107716 |
| *Neisseria gonorrhoeae* | Ne.go | Betaproteobacteria | NC_022240.1 | dRNA-seq | 919 | 25143534 |
| *Bordetella pertussis* | Bo.pe | Betaproteobacteria | NC_002929.2 | dRNA-seq | 593 | 29683387 |
| *Burkholderia cenocepacia* | Bu.ce | Betaproteobacteria | AM747720.1 | dRNA-seq | 1086 | 26462475 |
| *Xanthomonas campestris* | Xa.ca | Gammaproteobacteria | AM920689.1 | dRNA-seq | 1545 | 26975844 |
| *Acinetobacter baumannii* | Ac.ba | Gammaproteobacteria | CP000521.1 | dRNA-seq | 864 | 27806702 |
| *Pseudomonas aeruginosa* | Ps.ae | Gammaproteobacteria | NC_008463.1 | dRNA-seq | 2117 | 23028334 |
| *Pseudomonas putida* | Ps.pu | Gammaproteobacteria | NC_002947.3 | dRNA-seq | 1142 | 27111755 |
| *Shewanella oneidensis* | Sh.on | Gammaproteobacteria | NC_004347.2 | Multiple† | 1802 | 24987095 |
| *Vibrio cholerae* | Vi.ch | Gammaproteobacteria | NC_002505.1 | dRNA-Seq | 1283 | 25646441 |
| *Klebsiella aerogenes* | Kl.ae | Gammaproteobacteria | NC_015663.1 | dRNA-seq | 468 | 27806702 |
| *Salmonella enterica* Typhimurium | Sa.ty | Gammaproteobacteria | FQ312003.1 | dRNA-seq | 1120 | 22538806 |
| *Shigella flexneri* | Sh.fl | Gammaproteobacteria | CP037923.1 | dRNA-seq | 2051 | 32252626 |
| *Escherichia coli* | Es.co | Gammaproteobacteria | NC_000913.2 | dRNA-seq | 1865 | 25266388 |

\*TSS inferred based on the RNA-seq read coverage
†Tiling microarray and 5'-end RNA sequencing
‡Candidate phyla radiation

## Results

### Determining the-35 and-10 sequence–function relationship for model construction

We applied sort-seq to determine the sequence–function relationship of the-35 and-10 elements, using green fluorescent protein (GFP) expression of *E. coli* cells to report promoter strength^34^. A promoter library (PL) comprising all possible-35 and-10 sequence variants (4¹² = 16,777,216) was synthesized in the *E. coli* P*_purR_* promoter context (Fig. 2a). Through fluorescence-activated cell sorting (FACS) and deep sequencing, two sort-seq replicates collectively detected 11,670,734 promoter variants (69.6%; Fig. 2b). Among these, we retained 1,022,018 variants that passed a data quality filter (≥ 20 reads and present in at least two sort-seq ranks) for downstream analysis and model fitting. Measurements from the two replicates were highly correlated (Pearson’s *r* = 0.904; Fig. 2c), and promoter strength ranged from 2 to 1,115 relative fluorescence units (RFU; Fig. 2d), with 90% of variants ≤ 5.8 RFU.

We constructed a model based on an established statistical thermodynamics framework^21,35,36^. It assumes that a promoter exists in either the unbound or RNAP-bound state, with promoter strength proportional to the RNAP-bound probability (Fig. 2e). The relative probability of each state is determined by the Boltzmann factor e*^-βΔεRNAP^*, where *β* is a system constant and *−Δε_RNAP_* denotes the free energy difference between the RNAP-bound and unbound states. The *−Δε_RNAP_* is the sum of the free energy contributed by the-35 and-10 sequences and the spacer length.

Fitting the model to sort-seq determined promoter strength estimated these free energy parameters with the mean absolute error as 0.414 (Fig. 2f,g). Results indicate that the consensus - 35 and-10 sequences and the 17-bp spacer length conferred the strongest RNAP–promoter binding, consistent with the current knowledge (see Methods for details).

### Model predicts promoter architecture across diverse bacteria

We applied our model trained on the synthetic PL library to identify the-35 and-10 elements in native promoters across 49 species. Under a 15–19 bp spacer length constraint, our model scanned for potential-35 and-10 elements that maximized −Δε*_RNAP_* within the promoter region preceding each experimentally mapped TSS (Fig. 1c). To evaluate model performance in discerning promoter architecture in each species, we applied it to random genomic sequences and promoter-shuffled sequences, the latter retaining base frequencies but not order. Differences between the −Δε*_RNAP_* distributions of these sequences and native promoters were quantified by Kolmogorov-Smirnov statistics, with examples shown in Fig. 3a. Overall, the model distinguished native promoters from random genomic sequences (*D_g_*) better than from promoter-shuffled sequences (*D_s_*, Fig. 3b and Extended Data Fig. 2a). A cutoff of *D_g_*, *D_s_* < 0.2 was applied to exclude *Phytoplasma sp.* (Ph.sp), *Pseudomonas putida* (Ps.pu), and *Shigella flexneri* (Sh.fl) from further analysis (marked red in Figs. 3, 4, and 6; species acronyms listed in Table 1). Poor model prediction for these species likely stemmed from the data quality of the original TSS mapping experiments: (1) their TSS did not exhibit the purine bias observed in most bacteria^38^, as indicated by the *P*-values of the NRN motif in Fig. 4a; (2) the 5′ untranslated regions of their mRNA tended to be longer, whereas most bacteria had the length distribution peaked at 20−49 bp (Extended Data Fig. 2e); and (3) their RNAP, both at the promoter-contacting residues and at the whole-protein level, closely resembled those of their sister species (Figs. 1b and 7a and Extended Data Figs. 1 and 5a).

**Fig. 3.**
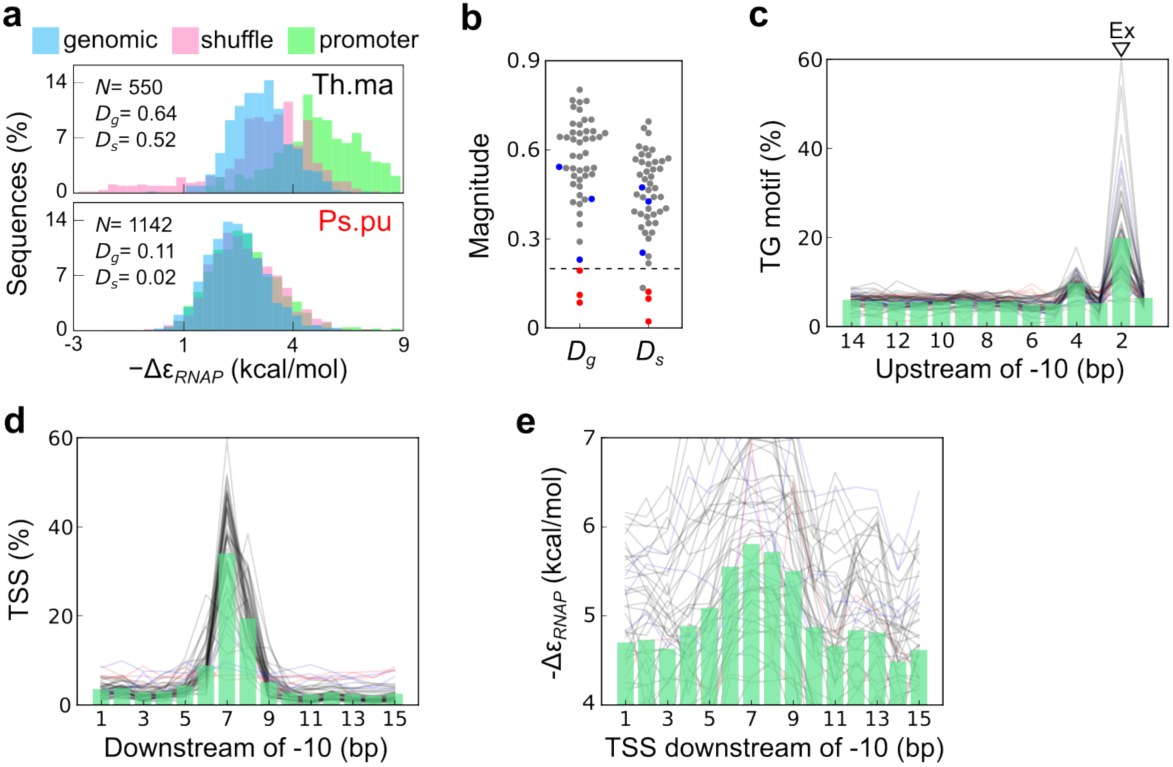
Model prediction and intrinsic properties of bacterial promoters.**a**, Model-predicted RNAP binding energy (−Δε*_RNAP_*) for *N* random genomic, promoter-shuffled, and native promoter sequences in distinguishable (e.g., *T. maritima*, Th.ma) and indistinguishable (e.g., *P. putida*, Ps.pu) cases. Differences between the −Δε*_RNAP_* distributions of native promoters and those of random genomic (*D_g_*) and promoter-shuffled (*D_s_*) sequences are quantified as Kolmogorov-Smirnov statistics. **b**, Summary of *D_g_* and *D_s_* values across the 49 species. The dashed line indicates a cutoff magnitude of 0.2. **c**, Sliding window analysis showing the frequency and the 3′ end location of the TG motif relative to the-10 element. **d**, Distribution of TSS relative to the-10 element. **e**, Correspondence between −Δε*_RNAP_* and the TSS position relative to the-10 element. In **a**-**e**, species with *D_g_*, *D_s_* < 0.2 or with greater variation in-10-to-TSS distance (standard deviation > 3.6 bp) are marked in red and blue, respectively. In **c**-**e**, individual species are represented by curves, while histograms show the mean distributions across the 49 species.

**Fig. 4.**
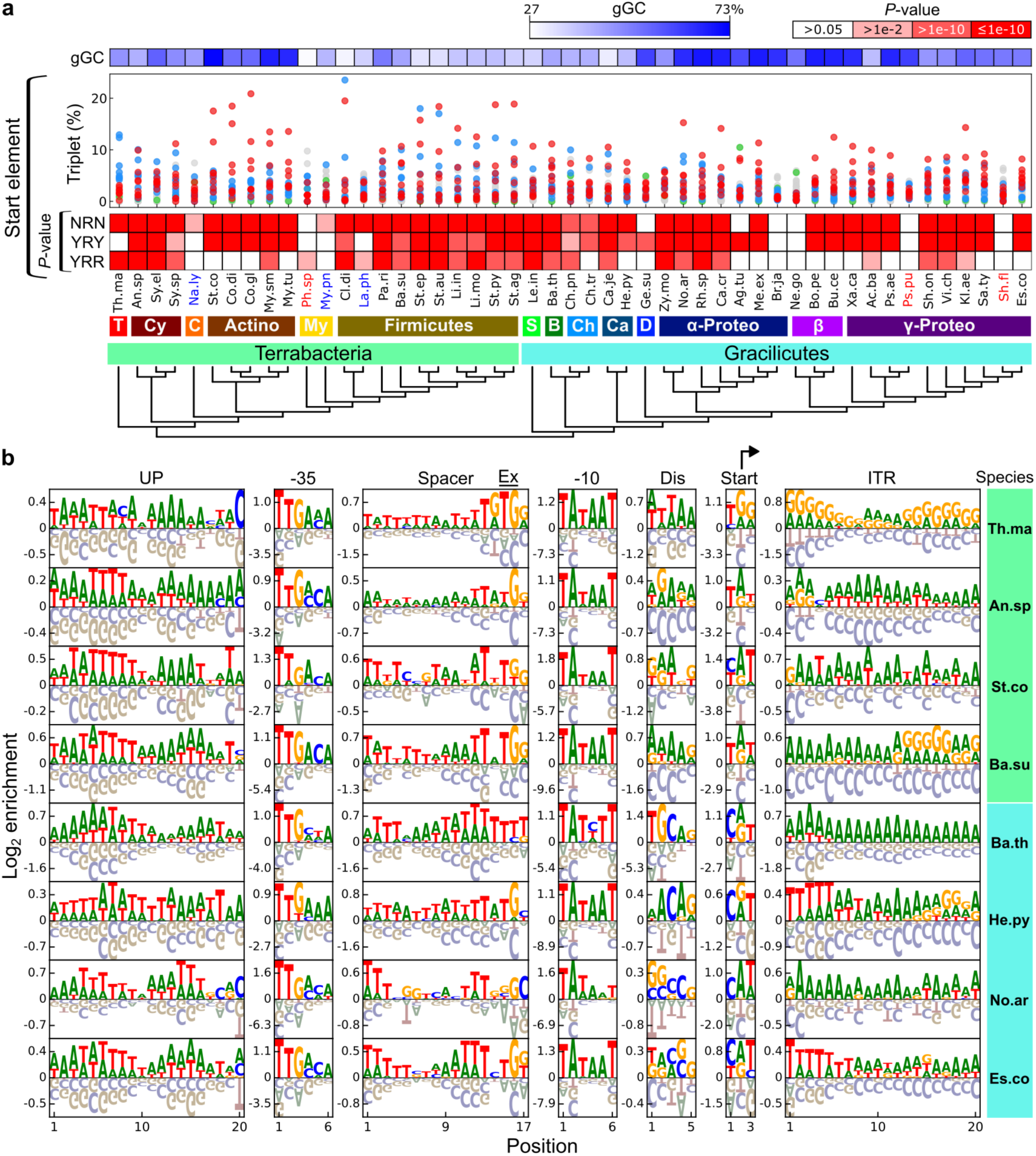
Characteristics of TSS, promoter elements, ITR, and GC content. **a**, Top to bottom: genomic GC content (gGC), proportions of all 64 triplets in the start element, statistical significances (*P*-value) of the TSS purine bias (i.e., NRN motif) and YRY and YRR motif frequencies conditioned upon TSS being purine, and the phylogenetic relationship of 49 bacteria^56^. Color bars in the top right indicate heatmap scales. Triplets belonging to YRY, YRR, or TTT motifs are indicated by red, blue, and green dots, respectively. Species with *D_g_*, *D_s_* < 0.2 or greater-10-to-TSS distance variation are marked in red and blue, respectively. T: Thermotogota, Cy: Cyanobacteria, C: Candidate Phyla Radiation, Actino: Actinobacteriota, My: Mycoplasmatota, S: Spirochaetota, B: Bacteroidota, Ch: Chlamydiota, Ca: Campylobacterota, D: Desulfobacterota, α-Proteo: Alphaproteobacteria, β: Betaproteobacteria, γ-Proteo: Gammaproteobacteria. **b**, Sequence enrichment in representative species. TSS is marked by a bended arrow. Graphic representation of positive and negative enrichment is rescaled for uniform visualization. Full enrichment analysis is presented in Extended Data Fig. 3. In **a**,**b**, species are shown as acronyms (full names listed in Table 1).

Trained on empirical data from *E. coli*, model predictions across species revealed that native promoters predominantly feature 17-bp spacer elements and-35 and-10 sequences resembling the consensus (Extended Data Figs. 2b and 3b). In addition, three properties—unique to native promoters yet not targeted by the model—support its translatability for mapping promoter architecture in phylogenetically diverse bacteria: (1) higher frequencies of TG and TRTG motifs at the Ex location (Fig. 3c and Extended Data Fig. 2c); (2) TSS was centered 7 bp downstream of model-predicted-10 elements (Fig. 3d and Extended Data Fig. 2d), consistent with empirical findings in *E. coli*^37^; (3) −Δε*_RNAP_* increased when the model predicted-10 elements to be 7 bp upstream of TSS (Fig. 3e and Extended Data Fig. 2d). Notably, the distance between TSS and model-predicted-10 elements was more variable in *Nanosynbacter lyticus* (Na.ly), *Mycoplasma pneumoniae* (My.pn), and *Lachnoclostridium phytofermentans* (La.ph) (standard deviations > 3.6 bp; marked in blue in Figs. 3, 4, and 6). Such variation may reflect either species-specific characteristics or reduced TSS-mapping accuracy owing to less common technologies (Table 1).

### Identifying a broadly conserved promoter element**—**start

Model mapping of the-35 and-10 elements in native promoters separated the length-variable UP, spacer, and discriminator elements. Based on these boundaries, we aligned each promoter element according to its length to reveal sequence enrichment, defined as the log-odds ratios of base frequencies relative to genomic backgrounds (Fig. 4b, Extended Data Fig. 3, and Supplementary Data 1). We chose this measure over the information content because it enhanced signal contrast by offsetting genomic composition bias (Methods).

Intriguingly, while aligning experimentally mapped TSS present in each species, we unveiled a conserved 3-bp motif, pyrimidine-purine-pyrimidine (Y^-1^R^+1^Y^+2^), centered on TSS (position +1). Among all 64 triplet bases, YRY was more frequent than the others in most bacteria examined (observations 30.6 ± 12.3% > null expectation 12.5%; Fig. 4a). Taking the genomic composition and purine bias of bacterial TSS into account^38^, YRY enrichment remained statistically significant (one-tailed binominal test: *P* < 0.05). Besides YRY, we noted the second abundant pyrimidine-purine-purine (Y^-1^R^+1^R^+2^) motif overrepresented in clade Terrabacteria (observations 28.7 ± 9.7% > null expectation 12.5%), particularly *Thermotoga maritima* (Th.ma; observation 56.0% > null expectation 12.5%), indicating lineage-specific divergence. In contrast, three species, *Geobacter sulfurreducens* (Ge.su), *Agrobacterium tumefaciens* (Ag.tu), and *Neisseria gonorrhoeae* (Ne.go), were enriched for the TTT motif (observations vs. null expectations as 4.9% vs. 0.7%, 10.5% vs. 0.8%, and 5.6% vs. 1.4%, respectively) and lacked TSS purine bias.

Among these 3-bp motifs, only YRY has been sporadically observed before in four species: *Acinetobacter baumannii* (Ac.ba), *Salmonella enterica Typhimurium* (Sa.ty), *Streptomyces coelicolor* (St.co), and *Mycobacterium tuberculosis* (My.tu)^39–42^.

Co-localization of the YRY motif with TSS suggests it role in TSS selection. To test the hypothesis and examine the effect of this motif on promoter strength, we analyzed data from another study that performed saturation mutagenesis to assess how *E. coli* RNAP positioned TSS in response to sequence variation 4–10 bp downstream of the-10 element (Fig. 5a)^37^. The authors observed the Y^-1^R^+1^ motif. By grouping all TSS based on their distance from the-10 element, we confirmed YRY enrichment and co-localization with TSS (Fig. 5b). Notably, co-localization occurred at 93.1% and 79.0% when the YRY motif was centered 7 or 8 bp downstream of the-10 element, respectively (Fig. 5c). Relative to YRY, the YRR motif had a similar but less prominent effect on TSS selection (86.2% and 67.4%, respectively). Apart from TSS selection, promoter strength increased significantly when TSS occurred 7 or 8 bp downstream of the-10 element, increasing further when YRY or YRR motifs co-localized (two-tailed Mann-Whitney U test: *P* < 10^-190^; Fig. 5d). In view of its functional significance in TSS selection and promoting transcription, broad conservation across the domain Bacteria, and divergence between the two major bacterial clades, Terrabacteria and Gracilicutes, we designated the 3-bp region centered on TSS as a new promoter element, ‘start.’

**Fig. 5.**
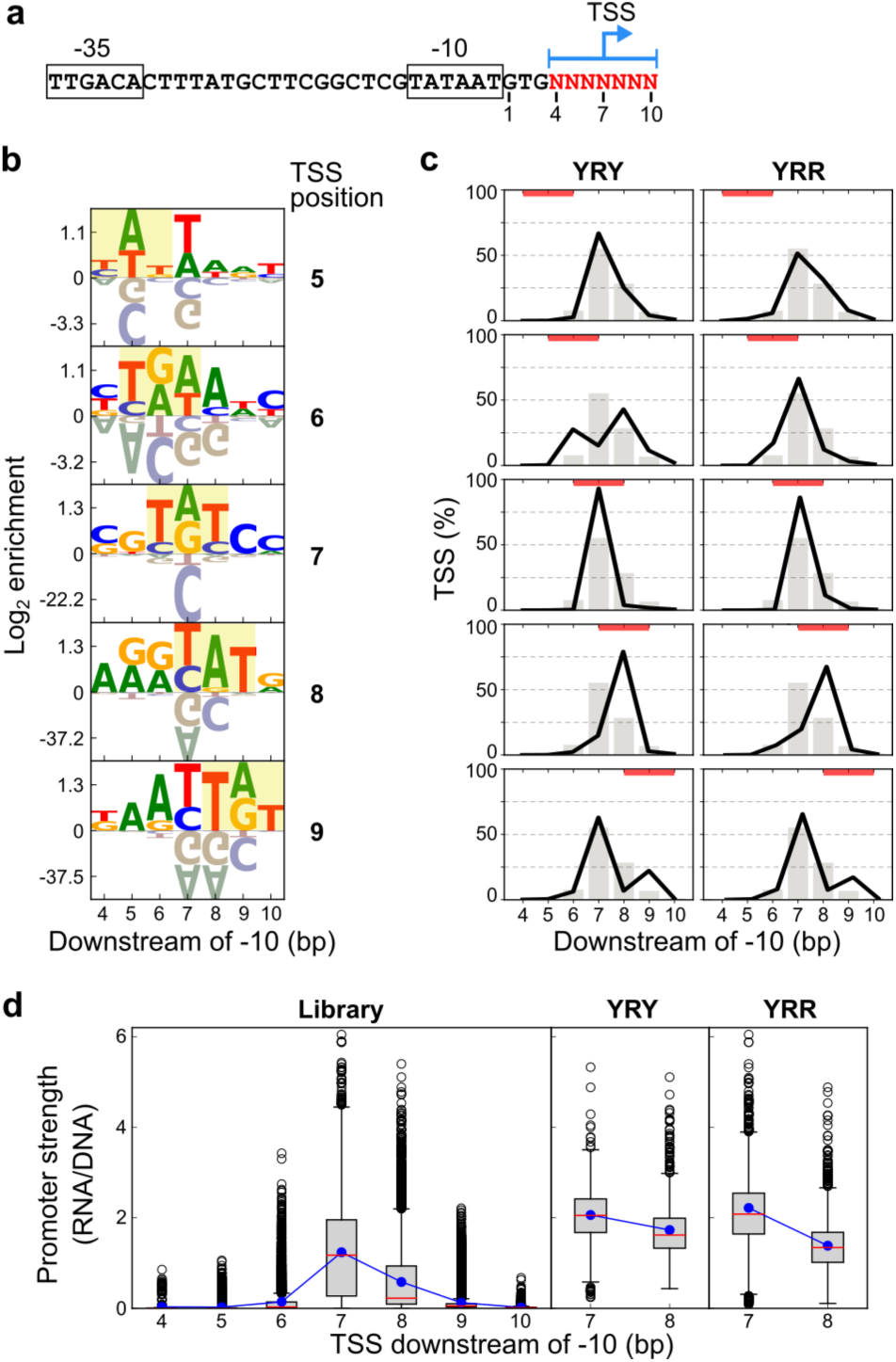
The start element influences TSS selection and promoter strength in *E. coli*. **a,** Promoter bearing a 7-bp randomized region (red) in a prior TSS mapping study^37^. **b,** Sequence enrichment in the 7-bp region for TSS occurring downstream of the-10 element. The 3-bp region centered on TSS is highlighted. Graphic representation for positive and negative enrichment is rescaled for uniformity. **c,** Influence of the location of YRY and YRR motifs (red bar) on TSS selection (black curve). The TSS distribution of the whole library (N = 16,212) is shown as the gray histogram. **d,** Library distribution of promoter strength with respect to the TSS location. Promoter strength is calculated as RNA reads divided by DNA reads (transcripts per promoter). The library subset, variants having YRY or YRR motifs co-localized with TSS = 7 or 8, are shown for comparison. Boxes, whiskers, blue dots, red lines, and empty circles indicate the interquartile range, 1.5-fold interquartile range, mean, median, and outliers, respectively, of the distribution.

### Variable promoter elements show broad conservation

Beyond the start element, we inspected enriched features in the other promoter elements. The-35 and-10 elements in most species were enriched for the consensus sequences (Fig. 4b and Extended Data Fig. 3b). In contrast, Alphaproteobacteria, including *Novosphingobium aromaticivorans* (No.ar), showed reduced-10 sequence conservation at its 3’ end. The-10 element of *Bacteroides thetaiotaomicron* (Ba.th) diverged even further, exhibiting a TATCTT hexamer followed by a TGC trimer extending into the Dis element. Both findings are consistent with previous reports^29,43^. The-35 element of *B. thetaiotaomicron* also evolved distinctly (TTGCTA), reflecting the significant divergence of its primary σ subunit (Fig. 1b and Extended Data Fig. 1)^44^. Recapitulating these deviations underscores the robustness of our model predictions.

In most species, the UP element exhibited alternating A-and T-tracts with a periodicity of roughly 10 bp (Fig. 4b). The wave-like pattern typically spanned 40 bp but extended up to 100 bp in Gammaproteobacteria (e.g., *E. coli* (Es.co) in Extended Data Fig. 3a). Additionally, the 3′ end of the UP element was often enriched for cytosine, suggesting its contact with R554 and R586 of σ^70^ is functionally important (Extended Data Fig. 1a)^45^. Regarding the spacer element, our model predicted its length to range mostly 16-18 bp, with 17 bp predominant across species (Extended Data Fig. 2b). Despite considerable sequence variation, the spacer element can be subdivided into four distinct regions in the 5′-to-3′ direction: an AT-tract (mainly thymine), a low-signal region with slight GC enrichment (e.g., *N. aromaticivorans*, *S. coelicolor*, and *Rhodobacter sphaeroides* (Rh.sp)), another AT-tract, and a G-rich region including the Ex element (Fig. 4b and Extended Data Fig. 3c). Among them, the AT-tract and the G-rich region proximal to the-10 element have been demonstrated to promote transcription in *E. coli* and *Mycolicibacterium smegmatis* (My.sm), respectively^4,5,46–48^. In contrast, the AT-tract proximal to the-35 element has been noted in *E. coli*^4^, while the adjacent 3′ GC-enriched region has not been reported in prior studies.

Similar to the start element, the Dis element and initial transcribed region (ITR) of mRNA (defined as the regions directly upstream and downstream of the start element, respectively, to avoid overlap) displayed lineage specificity. The Dis element was AG-or AT-rich in Terrabacteria but highly variable in Gracilicutes (Fig. 4b and Extended Data Fig. 3d). ITR was AG-rich in Firmicutes (e.g., *Bacillus subtilis* (Ba.su)) and *T. maritima*, AT-rich in Actinobacteriota (e.g., *S. coelicolor*), and frequently contained a 5′ T-tract in Gracilicutes (e.g., *Helicobacter pylori* (He.py); Fig. 4b and Extended Data Fig. 3e). To systematically assess the sequence conservation of each element within and between species, we applied two quantitative measures.

Intra-species conservation was estimated using Kullback-Leibler (KL) divergence, which gauged the information contrast between an element and its genomic background (Methods). As expected, the-10 and-35 elements ranked as the top two conserved elements across species (Fig. 6a). Notably, the start element ranked third, reaffirming its functional and evolutionary significance. Inter-species conservation was quantified as the sequence enrichment similarity of an element between two species, defined by the Spearman’s correlation of enrichment scores (e.g., the-10 element has 24 enrichment scores corresponding to four bases at six positions; Fig. 6b and Extended Data Fig. 4). Consistent with aforementioned observations, the-35 and-10 elements were highly conserved across species, except *B. thetaiotaomicron*. The UP, spacer, and Ex elements were moderately conserved, except *Synechocystis* sp. (Sy.sp), *R. sphaeroides, Leptospira interrogans* (Le.in)*, Mycoplasma pneumoniae* (My.pn), and *B. thetaiotaomicron*.

**Fig. 6.**
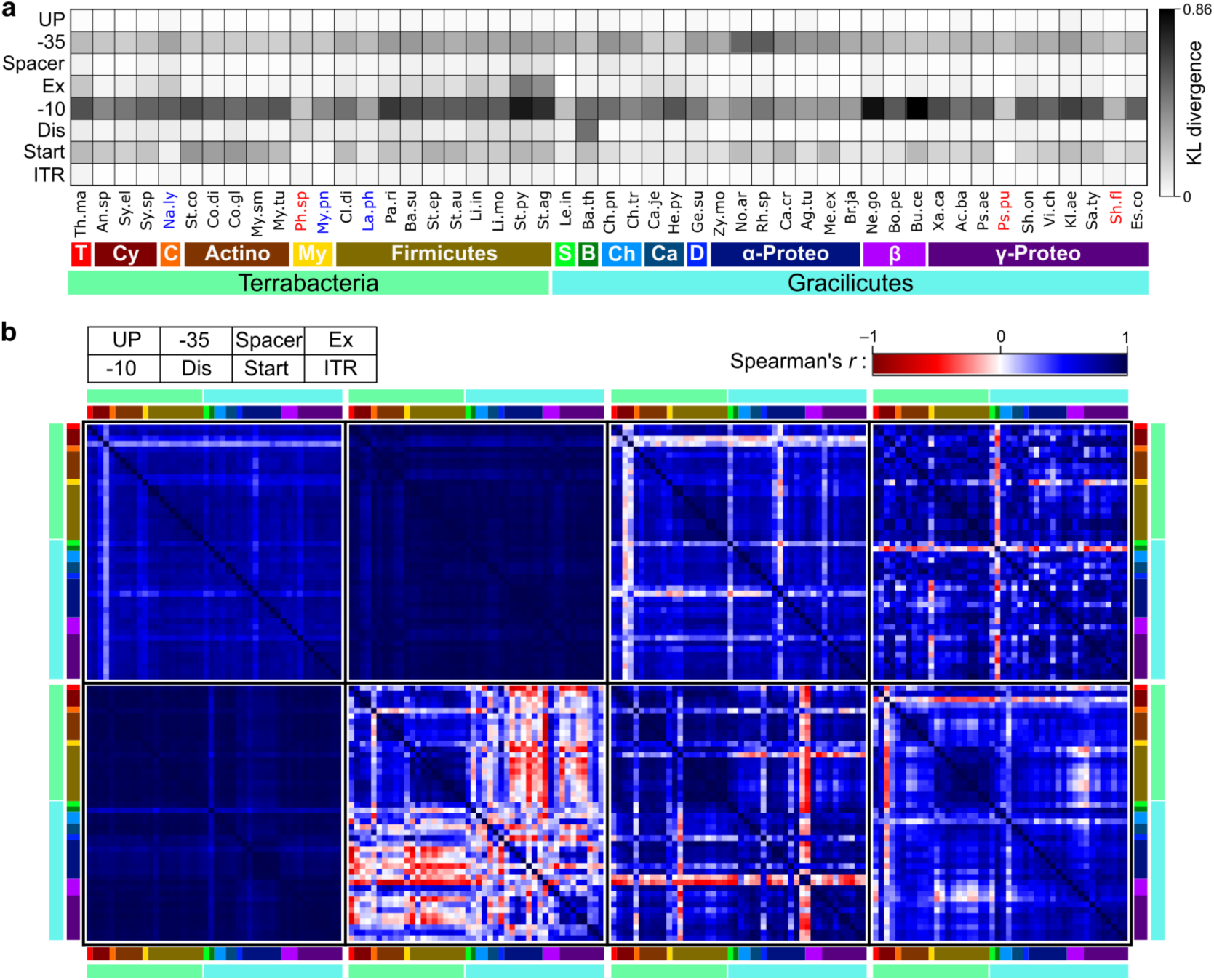
Conservation of promoter elements and ITR within and between species. **a**, Intra-species conservation quantified by KL divergence per bp. **b**, Inter-species conservation measured as the Spearman’s correlation of enriched sequence features. Species having *D_g_*, *D_s_* < 0.2 (i.e., Sh.fl, Ps,pu, Ph.sp) are excluded. Lineages are distinguished by colors shown in **a**. In **a**,**b**, analyses consider the dominant length type of the Dis (5-bp) and spacer elements (17-bp, excluding the 2-bp Ex region), the 40-bp UP element region upstream of the-35 element, and the 40-bp ITR region downstream of the start element. The abbreviations of species and lineage names follow Fig. 4 and Table 1.

**Fig. 7.**
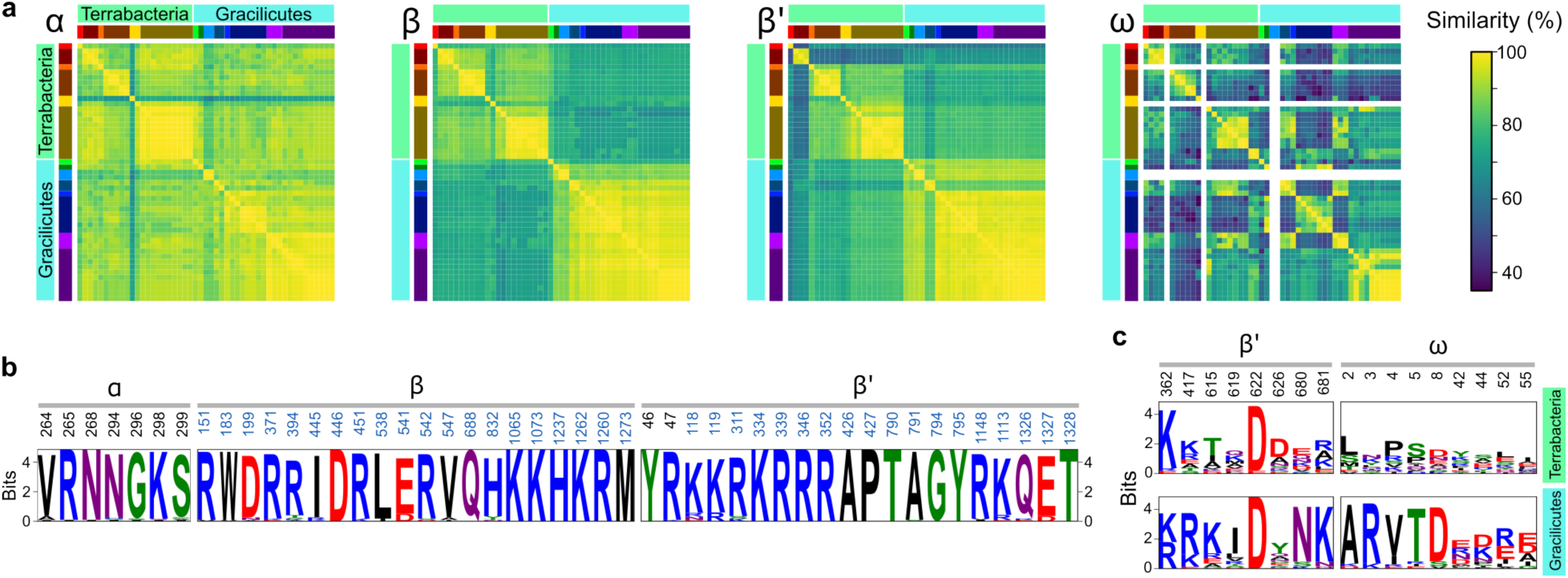
Sequence conservation of the RNAP α, β, β’, and ω subunits. **a**, Pairwise protein sequence similarity of the α, β, β’, and ω subunits among the 49 species. Species are ordered and lineages are distinguished by colors, as in Fig. 4a. The ω subunit could not be identified in *N. lyticus*, *M. pneumoniae*, *C. pneumoniae*, and *C. trachomatis* based on genome annotation or BLAST searches, and is hence left blank. **b**, Conservation of promoter-contacting residues quantified by information content^13,45,61,62^. Residue positions correspond to *E. coli* homologues, with positions contacting the Dis and start elements and ITR marked in blue. **c**, Conservation of ppGpp-contacting residues in the β’ and ω subunits in Terrabacteria and Gracilicutes. Sequence alignments for **b**,**c** are shown in Extended Data Fig. 5.

Overall, promoter elements and ITR were more conserved within Firmicutes and Actinobacteriota than the other lineages (Fig. 6b). Relative to the-10 element and those further upstream, the Dis and start elements and ITR showed greater differentiation between the two major bacterial clades. In particular, the Dis and start elements were more conserved within Terrabacteria than within Gracilicutes. Despite these between-clade differences, residues in the σ^70^, β, and β′ subunits that contact the Dis and start elements and ITR remain broadly conserved (Figs. 1a and 7b, Extended Data Figs. 1a and 5a). Given that these subunits show between-clade divergence at the whole-protein level (Figs. 1b and 7a), RNAP residues beyond the promoter-contacting sites may contribute to the observed clade distinction in promoter elements and ITR.

### Positive selection shapes broad spacer sequence conservation

To investigate the importance of sequence conservation in the spacer element, we performed sort-seq to map its sequence–function landscapes in *E. coli*. We synthesized saturation mutagenesis libraries, SL16, SL17, and SL18, for 16-, 17-, and 18-bp spacer elements, respectively, paired with distinct-35 and-10 elements to reduce potential context dependence (Fig. 8a). There, the-35 and-10 elements were designed to differ 1–3 bp from the consensus sequences to mitigate their dominant influence on transcription initiation, as revealed by prior studies^49,50^. From 3,855,981, 6,984,696, and 12,242,605 detected variants, we retained reliable measurements, divided each library into five groups based on promoter strength, and computed each group’s sequence enrichment relative to the whole library (Fig. 8b and Extended Data Fig. 6). Despite differences in spacer length and promoter context, the three libraries revealed parallel characteristics from low to high promoter strength (Groups 1 to 5 in Fig. 8c): (1) the spacer element contained four distinct regions, mirroring those observed across species (Fig. 4b); (2) enriched sequences in Group 5 resembled those in natural spacer elements, except for an apparent lack of the signature TG motif of the Ex element. Notably, the TG motif frequency increased steeply with promoter strength (Fig. 8d), indicating its positive contribution to transcription. Underrepresentation of the TG motif was therefore an artifact of enrichment analysis, masked by the positive enrichment of guanine (Fig. 8c,d).

**Fig. 8.**
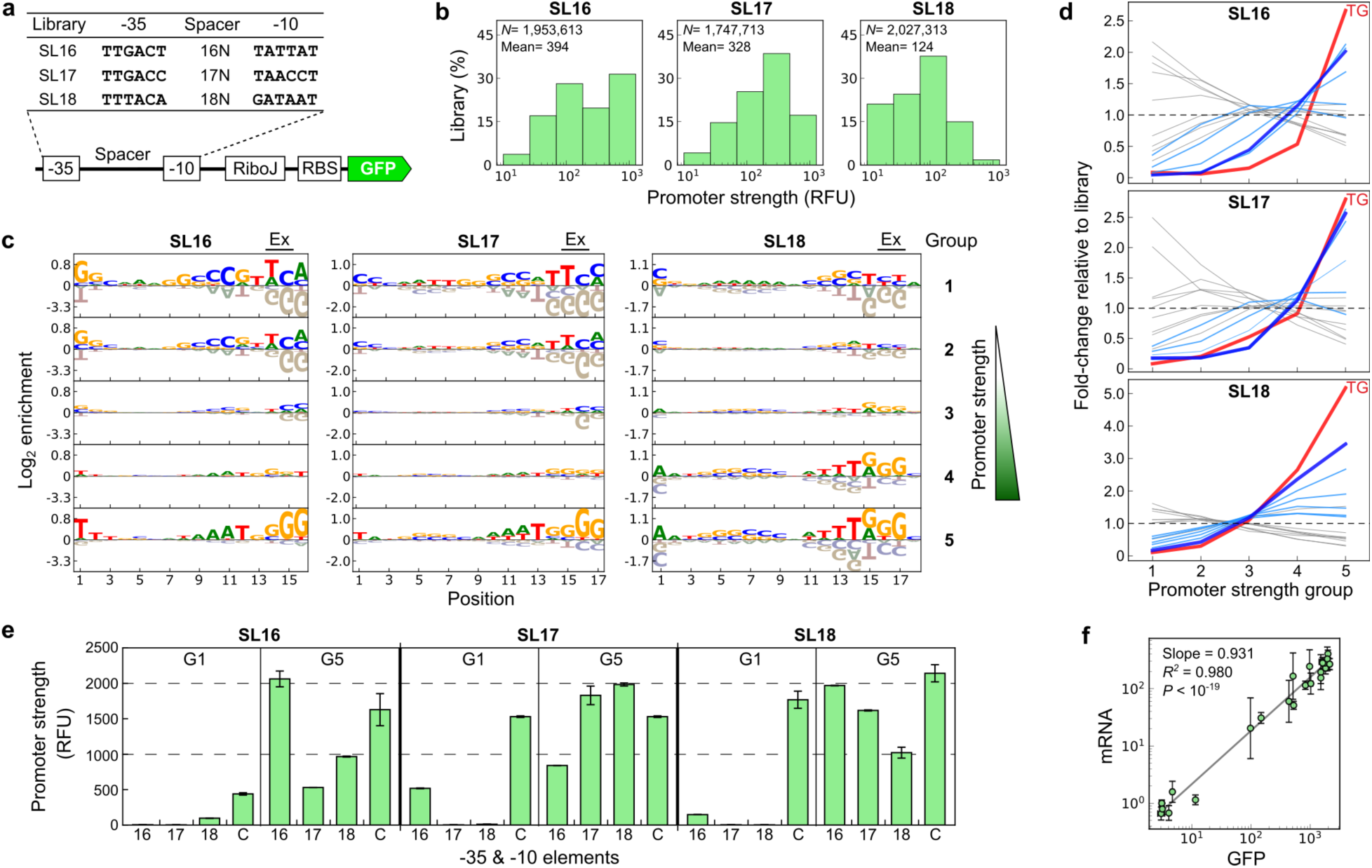
Determining the spacer sequence–function relationship. **a**, Promoter backgrounds of the SL16, SL17, and SL18 libraries. **b**, Library distribution of promoter strength in *E. coli*. RFU, relative fluorescence unit. **c**, Enriched sequences in the spacer element with respect to promoter strength. Spacer variants are grouped based on the binning in **b**. Graphic representation for positive and negative enrichment is rescaled for uniformity. **d**, Fold-changes in the frequencies of the 16 2-bp motifs in the Ex region relative to the library (dashed line) across groups. The trends of ‘TG,’ ‘GG,’ and the remaining G-bearing motifs are marked in red, deep blue, and light blue, respectively. **e**, Transcriptional impact of the spacer element across promoter contexts. Promoters consist of the consensus or library-35 and-10 sequences shown in **a** (denoted as C, 16, 17, 18), paired with positively enriched spacer sequences from Groups 1 or 5 (denoted as G1, G5) of each spacer library shown in **c**. Results are shown as the means and standard deviations of three independent measurements. **f,** Linear regression of mRNA fold-changes against GFP levels (i.e., promoter strength) for the 24 promoter variants shown in **e**. The lowest-expressing variant (SL17-35/-10 sequences paired with its G1 spacer) serves as the reference for calculating mRNA fold-changes. Dots and error bars represent means and standard deviations from at least three independent measurements.

To verify the context independence of the above findings, we characterized synthetic promoters in which the enriched spacer sequences of Groups 1 and 5 (denoted as G1 and G5, respectively) from each spacer library were paired with either consensus or the three library-35/-10 sequences (denoted as 16, 17, 18, and C, respectively in Fig. 8e). Across SL16, SL17, SL18, G5 spacers conferred significantly higher promoter strength than G1 spacers when paired with the four sets of-35/-10 sequences (*P* < 0.003, paired *t*-test). The only exception was observed in SL17—its G1 and G5 spacers conferred similar promoter strength when paired with the consensus-35/-10 sequence (*P* > 0.05, *t*-test), suggesting that the consensus-35/-10 sequence, under the optimal 17-bp spacer length, dominates and saturates RNAP–promoter binding potency. Remarkably, the G1 and G5 spacers from SL16 exhibited a > 600-fold promoter strength difference when paired with the SL16-35/-10 sequences, underscoring the transcriptional impact of the spacer sequence composition. These findings, validated by both GFP and mRNA expression (Fig. 8f), confirm the functional importance of the four-region organization of the spacer element and suggest that positive selection shapes its broad conservation in nature.

### Diversifying evolution of the Dis element driven by growth rate regulation

To probe the function of variable Dis elements in Gracilicutes, we performed saturation mutagenesis and sort-seq to characterize the 8-bp region directly downstream of the-10 element in *E. coli* (Fig. 9a). Promoter strength in this DL library spanned a 240-fold range. Based on promoter strength, we divided the DL variants into five groups to examine their sequence enrichment (Fig. 9c). From low to high promoter strength (Groups 1 to 5), enriched features shifted from GC-rich to C-poor sequences, with the 5‘ 3-bp region showing stronger enrichment (Fig. 9d). The GC-rich pattern mirrored the 8-bp Dis element in the ribosomal promoter *rrnB* P1 (P1*_rrnB_*; Fig. 9b)^15^. Paradoxically, the exact sequence was among the lowest-expression variants in the DL library (24^th^ in 58,065; Fig. 9c), while P1*_rrnB_* was among the strongest promoters in *E. coli*^51^.

**Fig. 9.**
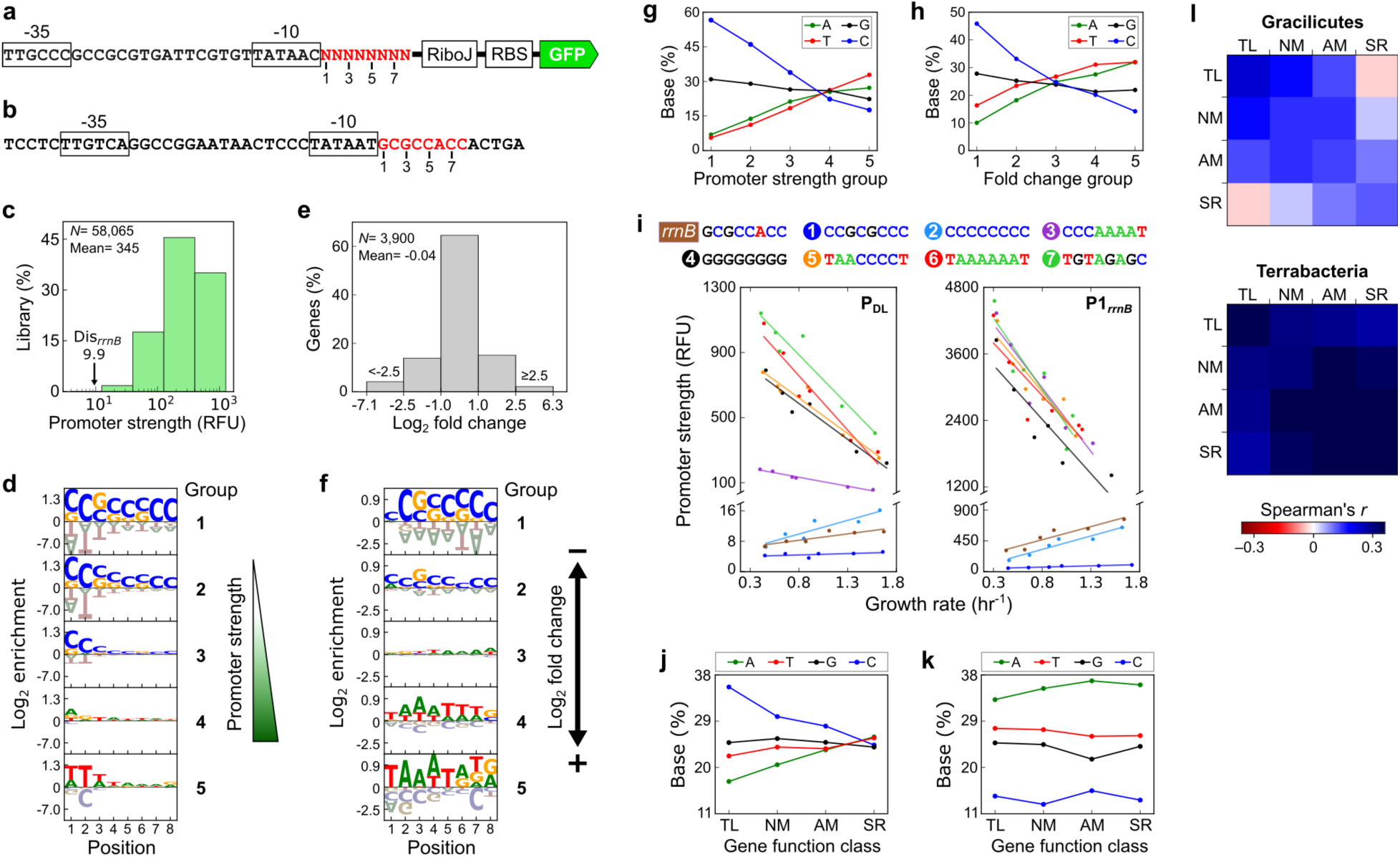
Determining the Dis sequence–function relationship. **a**,**b**, DL library promoter background (**a**) and P1*_rrnB_* promoter (**b**), with the 8-bp Dis marked in red. **c**, DL library distribution of promoter strength in *E. coli*. Dis*_rrnB_*, variant bearing the P1*_rrnB_* Dis in **b**. **d**,**g**, Sequence enrichment (**d**) and base composition (**g**) of DL variants with respect to promoter strength. Variants are divided into five groups based on the binning in (**c**). **e**, Distribution of the mRNA fold-changes of *E. coli* genes in response to ppGpp elevation^12^. **f**,**h**, Sequence enrichment (**f**) and base composition (**h**) of *E. coli* native Dis with respect to ppGpp-induced transcriptional changes^12^. Native Dis is grouped based on the binning in (**e**). **i**, Promoter strength of DL variants (top) in the library (P_DL_) or P1*_rrnB_* promoter contexts across growth rates in *E. coil*. Dots and lines represent data and linear regression curves, respectively. All slopes are statistically significant (R^2^ > 0.75, *P* < 0.03), except DL_1_ variant in the P_DL_ context. **j**,**k**, Base composition of Dis sequences associated with genes involved in translation (TL), nucleotide metabolism (NM), amino acid metabolism (AM), and signal response (SR) in Gracilicutes (**j**) and Terrabacteria (**k**). **l**, Similarity (Spearman’s correlation) of Dis sequence enrichment across TL, NM, AM, and SR genes. Shown here are the mean similarities within and between gene function classes across species, based on Extended Data Fig. 7g. Species are excluded if they have *D_g_*, *D_s_* < 0.2 or < 10 Dis sequences in a given class. In **d**,**f**, graphic representation for positive and negative enrichment is rescaled for uniformity.

Why would a strong promoter bear a strength-compromising component? A prior study demonstrated that the GC-rich Dis element made P1*_rrnB_* transcription scale positively with growth rate^15^. This growth rate-dependent regulation results from increasing production of the stringent response signal ppGpp under slow growth, which allosterically regulates RNAP to reduce P1*_rrnB_* transcription^52–54^. Another study further reported changes in 35.4% of the *E. coli* transcriptome due to exogenous ppGpp overproduction (Fig. 9e)^12^. Among these changes, genes within the same functional class showed correlated responses, for example: transcription of translation-and nucleotide metabolism-related genes decreased significantly, whereas transcription of amino acid metabolism-and signal response-related genes increased strongly.

Coincidentally, Dis sequences in native *E. coli* promoters whose transcription responded negatively and positively to ppGpp (Groups 1 to 5 in Fig. 9f) resembled the low-and high-expression variants in the DL library (Groups 1 to 5 in Fig. 9d), respectively, despite the DL library being assayed in a nutrient-rich and ppGpp-poor condition. Examining the base composition of DL variants and native Dis elements across groups further highlighted their similar design principles (Fig. 9g,h).

Based on above findings, we hypothesized that high Dis variation within Gracilicutes arose in part from diversifying evolution, enabling cells to fine-tune individual promoters to coordinate the gene expression of each functional class across growth rates. If so, we would expect: (1) growth rate dependency conferred by Dis elements to be modular; (2) ppGpp-contacting residues on RNAP to be conserved; and (3) genes within the same functional class to share similar Dis sequence features. To test modularity, we selected eight DL variants from low-to high-expression and examined their effects on promoter strength in the P1*_rrnB_* and DL promoter contexts (P_DL_) in *E. coli* across growth rates (0.3-1.7 hr^-1^; Fig. 9i). Despite the distinct expression ranges of P_DL_ and P1*_rrnB_*, seven of eight DL variants exerted similar effects on expression and growth rate dependency in both promoter contexts. Moreover, low-and high-expression DL variants consistently led to positive and negative correlations, respectively, between promoter strength and growth rate. Next, we examined ppGpp-contacting residues on the β’, and ω subunits of RNAP and found higher conservation in Gracilicutes than Terrabacteria (Fig 7c and Extended Data Fig. 5b)^53^.

Finally, we examined native Dis sequences of *E. coli* genes classified under translation, nucleotide metabolism, amino acid metabolism, and signal response, according to KEGG Orthology^55^. Compared with genome-wide Dis sequences, those of the four gene function classes shifted from being overall C-rich to diverse and C-reduced in Gracilicutes (Fig. 9j), echoing *E. coli* Dis design principles (Fig. 6g,h). In contrast, Dis sequences in Terrabacteria were AG-or AT-rich regardless of gene function classes (Fig. 9k and Extended Data Fig. 7a-e). Examining the similarity of Dis sequences across the four gene function classes, defined by the Spearman’s correlation of enrichment scores (i.e., 32 scores corresponding to four bases at eight positions), further revealed the higher Dis sequence conservation among translation-related genes and the contrasting Dis features between translation-and signal response-related genes in Gracilicutes (Fig. 9l and Extended Data Fig. 7f,g). Together, these findings support an evolutionary scenario—growth rate-dependent regulation in Gracilicutes has imposed functional constraints on RNAP and diversifying selection on the Dis element, resulting in regulatory divergence of RNAP and core promoters between the two major bacterial clades.

## Discussion

Variation in the sequence and length of promoter elements makes computational identification and functional characterization of promoters challenging. To address this, we developed a simple model capable of mapping core promoter organization across phylogenetically diverse bacteria using only the-35 and-10 sequences and spacer length. Though the model-predicted −Δε*_RNAP_* values of native promoters were generally higher than those of random sequences in most species, their distributions overlapped substantially (Fig. 3a and Extended Data Fig. 2a). In addition, model prediction currently requires experimentally mapped TSS to define promoter regions. While the results highlight the potential of our model for genome-wide and cross-domain promoter mapping, its limitations point to the need for further refinement. To this end, future model improvement can incorporate the conserved features of the other promoter elements, as revealed in Fig. 4b. In addition to advancing native promoter identification, our experimentally determined sequence–function relationships for the-35, spacer,-10, and Dis elements provide invaluable data for training models aimed at designing promoter sequences with precise expression levels for synthetic biology applications. Among these, the spacer element alone can modulate promoter strength by up to 600-fold (Fig. 8e), underscoring its significance in regulating transcriptional initiation. Lineage-and gene function-specific traits, such as Dis sequence divergence (Fig. 9), offer further guidance for engineering promoters beyond model organisms and crafting promoters with the growth rate-tunable property.

In parallel to bioinformatic and bioengineering applications, this study yields critical insights into the evolution of transcription mechanisms. Our findings emphasize the marked differences in RNAP subunits and core promoter elements (Dis and start) between Terrabacteria and Gracilicutes (Figs. 1b, 4a, 6b, 7a, 7c, 9j-l), underscoring the divergence of transcriptional regulation between the two major bacterial clades beyond their distinct cell wall organization^56,57^. Among these divergent features, gene function-dependent Dis similarity (Fig. 9j,l), together with systems-level modulation of cellular RNA polymerase and ribosome concentrations^58,59^, illustrate multilayered gene regulation through which Gracilicutes orchestrate global gene expression in response to changing growth rates. Future molecular and biochemical studies could use synthetic DNA fragments bearing these clade-specific features, together with RNAP from diverse species or reconstructed through phylogenetic resurrection, to elucidate the molecular mechanisms underlying evolutionary transition and functional differentiation in RNAP–promoter interactions. Extending to the cross-domain context, identification of sequence and functional similarities between bacterial promoter elements, start (YRY) and-10 (TATAAT), and their archaeal and eukaryotic counterparts, initiator (YR) and TATA box (TATAAAA), strongly suggests a shared evolutionary origin of promoter architecture across the three domains of life. This core promoter organization is retained, despite significant structural divergence of core RNAP and recruitment of non-homologous subunits that recognize the-10 element (σ) in bacteria and the TATA box (TATA-binding protein) in the archaea/eukaryote lineage^32,60^.

## Methods

### Development of a biophysical model

Our model assumes that promoters exist in either the unbound or RNAP-bound state, with promoter strength proportional to the RNAP-bound probability. The state’s relative probability is determined by the Boltzmann factor *e^-βΔε^*, where *−Δε* denotes the free energy, *β = 1/RT*, gas constant *R* = 1.98e-3 kcal mol^-1^ K^-1^, and *T* = 310 K. The unbound-state free energy is set to zero as the ground condition. The free energy of the RNAP-bound state (*−Δε_RNAP_*) is the sum of the free energies of the-35 element (*−Δε_-35_*), spacer element (*−Δε_Spacer_*),-10 element (*−Δε_-10_*), and the P*_purR_* promoter context (*−Δε_0_*). The model assumes each base (*b*) at each position (*i*) in the - 35 and-10 elements contributes to free energies additively (i.e., *−Σ_i_ Δε_i_*(*b*)). It further assumes *−Δε_Spacer_*(*l*) varies with the spacer length (*l*) regardless of sequence^64^. During model fitting, the model scanned through each PL variant sequence, identified all potential-35 and-10 element pairs with spacer length as 16-18 bp, and selected the configuration that maximized *−Δε_RNAP_* as the promoter. The model assumes the transcription rate constant as *r_max_* in the bound state, whereas there exists basal measurement noise *r_min_* in the unbound state, with *r_min_* << *r_max_*. We estimated model parameters by minimizing the mean squared error using the Python SciPy and scikit-learn packages^65^. The PL library promoter strength distribution was divided into ten equal-sized bins, from which 1,000 promoter variants per bin were randomly sampled with replacement as the training datasets (Fig. 2d)^66^. Model fitting estimated *−Δε_0_* = *−*7.3, *r_min_* = 3.13, the free energies of the 16-bp (*−Δε_Spacer_*(16)) and 18-bp (*−Δε_Spacer_*(18)) spacer elements, and that of each base in the-35 and-10 elements (Fig. 2g). Due to the scarcity of high-expression variants, we set *rₘₐₓ* to 1000 RFU (i.e., maximum observed in the PL library) to represent the plateau of the model-fitted sigmoidal curve for stable parameter estimation (Fig. 2f). We tested this assumption by performing model training 30 times with *rₘₐₓ* as a free parameter, resulting in *rₘₐₓ*= 452–774 RFU. Despite this variation, the inferred −Δε*ᵢ*(*b*) and *−Δε_Spacer_*(*l*) correlated strongly with the *rₘₐₓ*-constrained estimates (Pearson’s *r* = 0.981–0.985). Similarly, to resolve parameter degeneracy that would arise due to independent estimation of *−Δεᵢ*, *−Δε_Spacer_*, and *−Δε₀*, we followed a previous modeling procedure to set *−Σ_b_ Δε_i_*(*b*), *−Δε_Spacer_*(17) = 0^24^. To evaluate parameter sensitivity to the reference point, we performed model training 30 times using the same dataset while varying *−Δε_Spacer_*(17). The resulting −Δε*ᵢ*(*b*) were highly correlated (Pearson’s *r* = 0.982–0.983). To expand the model prediction range for spacer length, we adopted *−Δε_Spacer_*(15) and *−Δε_Spacer_*(19) estimated by another study^64^. The two parameters were rescaled to adjust for the numerical difference, reflected by the fold difference of *−Δε_Spacer_*(16), *−Δε_Spacer_*(17), and *−Δε_Spacer_*(18) between the two studies.

### Model prediction and sequence analysis of promoter elements

We applied the biophysical model to predict promoter elements across the domain Bacteria. We collected the experimentally determined TSS information of 49 diverse species from the literature (Table 1). Among them, 47 TSS datasets were obtained by dRNA-seq or other direct detection methods, while two (*N. lyticus* and *M. pneumoniae*) were indirectly inferred based on the abrupt increase of the RNA-seq read coverage upstream of genes. Only primary TSS, the dominant TSS in a TSS cluster upstream of genes^67^, was chosen for model prediction. The model scanned the 46-bp region upstream of each TSS to predict the-35 and-10 elements based on the criteria that their hexamer composition and the spacer length (15-19 bp) maximized *−*Δε*_RNAP_*. To verify the model’s ability to identify true promoters, we created two negative control datasets with the sample size identical to the TSS dataset: one containing 46-bp random genomic sequences and the other containing promoter-shuffled sequences generated by permutating the 46-bp sequence upstream of each TSS. Model prediction of the-35 and-10 elements and identification of the start element (i.e., the 3-bp region centered upon TSS) defined ITR and the remaining promoter elements for each TSS (Supplementary Data 1). Sequence features in each promoter element and ITR were revealed by Log_2_ enrichment and KL divergence analysis. Let *p_i_* (*b*) and *q_i_* (*b*) denote the frequencies of base *b* ∈{A, T, G, C} at position *i* in a data subset and the whole dataset, respectively, where Σ*_b_ p_i_* (*b*) = Σ*_b_ q_i_* (*b*) = 1. The Log_2_ enrichment *E_i_* (*b*) was computed as *E_i_* (*b*) = log_2_ (*p_i_* (*b*)/*q_i_* (*b*)). The KL divergence *D_i_* of position *i* was calculated as *D_i_* = Σ*_b_ p_i_* (*b*) · ln (*p_i_* (*b*)/*q_i_* (*b*)).

### Sequence and phylogenetic analysis of RNAP subunits

We applied Biopython 1.81 to collect all coding sequences (CDS) of the 49 species from their genome files (Table 1)^65^. We then used RpoA, RpoB, RpoC, and RpoD of *E. coli* as the query sequences for BLAST, setting E-value =1e-5 as the cutoff to identify the homologous proteins of α, β, and β‘ subunits and σ^70^, respectively (Extended Data Figs. 1 and 5). BLAST with *E. coli* RpoA identified exactly one α subunit homolog per species. BLAST with *E. coli* RpoB identified one β subunit homolog per species, but the corresponding CDS in *H. pylori* additionally encoded the β‘ subunit at the C-terminal. Based on multiple sequence alignment, we divided this CDS into N-and C-terminal segments homologous to the β and β‘ subunits, respectively. We used the MUSCLE package in Jalview 2.11.3.3 with default settings to generate multiple sequence alignments, both here and subsequently^68,69^. BLAST with *E. coli* RpoC found two CDS in each of *Anabaena* sp., *Synechococcus elongatus*, and *Synechocystis* sp. as the cyanobacterial β‘ subunit has evolved into two smaller γ and β‘ subunits^70^. We identified their protein residues homologous to the full-length β‘ subunit by performing multiple sequence alignments with the β‘ subunits of the remaining 46 bacteria. Then, the homologous segments were concatenated to create a chimeric full-length β‘ subunit sequence for each cyanobacterium. BLAST with *E. coli* RpoD found 164 CDS showing significant sequence similarity in 49 bacteria. To distinguish the σ^70^ subunit from other paralogs in the same protein family, we constructed a maximum likelihood (ML) tree using the smart model selection (SMS) and fast likelihood-based method of PhyML 3.0^71,72^. Results revealed a 56-taxa clade that comprised single CDS from 40 species (e.g., *E. coli* RpoD) and 2-4 CDS from species belonging to Actinobacteriota (*S. coelicolor*, *Corynebacterium diphtheriae*, *Corynebacterium glutamicum*, *Mycolicibacterium smegmatis*, *Mycobacterium tuberculosis*) and Firmicutes (*L. phytofermentans*, *Clostridioides difficile*). As two species, *Phytoplasma* sp. and *Burkholderia cenocepacia*, had no CDS identified by BLAST due to missing annotation in their genome files, we separately retrieved the corresponding CDS (BAD04713.1 and CAR54775.1) from the NCBI GenBank and merged them with the 56-taxa clade, making the σ^70^ subunit collection. The σ^70^ subunit collection was subjected to multiple sequence alignment, followed by ML tree construction using SMS and the standard bootstrap analysis (n=1,000) of PhyML 3.0 (Extended Data Fig. 1)^71,72^. The ML tree was visualized by FigTree 1.4.4^73^. The pairwise sequence similarity of CDS was calculated using the PairwiseAligner function in the Bio.Align package of Biopython 1.81 using BLOSUM62 as the scoring matrix. Since the RNAP ω subunit is too short and variable to be reliably identified by BLAST, we retrieved its CDS primarily through finding annotations that contain keywords ‘omega’ or ‘rpoZ’ in genome files. We obtained single CDS of the ω subunit in 46 of 49 bacteria. We used *E. coli* RNAP ω subunit (RpoZ) as the BLAST query sequence but still could not find CDS with E-value ≤ 1e-5 in *N. lyticus*, *Chlamydia pneumoniae*, and *Chlamydia trachomatis*. The three species may lack the ω subunit since it is not essential. Multiple sequence alignment of the ω subunit was performed to examine its sequence conservation.

### Strain and growth medium

*E. coli* K-12 MG1655 was the host strain for experiments throughout this study. LB medium comprised 10 g NaCl (BioShop), 10 g bacterial tryptone (BD Bacto), and 5 g yeast extract (BD Bacto) in one liter of deionized water. LBK medium, LB medium supplemented with filter-sterilized kanamycin sulfate (final concentration as 50 mg/L, Thermo Fisher Scientific), was used for growing cells for plasmid extraction. LBKG medium, LBK medium supplemented with filter-sterilized glucose (final concentration as 10 g/L, Bionovas) to enhance growth and GFP expression, was used in growth assays or FACS experiments. One liter of M9 minimal medium comprised 5.98 g Na_2_HPO_4_ (AppliChem), 3 g KH_2_PO_4_ (AppliChem), 0.5 g NaCl, 0.8 g NH_4_Cl (AppliChem), 976.7 mL of deionized water, and the following components that were filter-sterilized separately and added immediately before use: 1 mL of 0.1 M CaCl_2_ (Hayashi Chemical), 2 mL of 1 M MgSO_4_ (Sigma-Aldrich), 0.2 mL of 185 mM FeCl_3_ (AppliChem), 0.3 mL of 1 mM thiamine hydrochloride (WS Simpson), 9.8 mL of M9 trace element solution, 1 mL of kanamycin sulfate (50 mg/mL), and the intended amount of carbon source and amino acid supplements. M9 trace element solution comprised 0.18 g ZnSO_4_·7H_2_O (J.T.Baker), 0.12 g CuCl_2_·2H_2_O (Sigma-Aldrich), 0.12 g MnSO_4_·H_2_O (J.T.Baker), and 0.18 g CoCl_2_·6H_2_O (Sigma-Aldrich) in 980 mL of deionized water. The final concentrations of glucose and glycerol (J.T.Baker) were 5 g/L and 2.2 g/L, respectively. The final concentrations of casamino acids (Bioshop) were 2 or 10 g/L (denoted as 1×CA and 5×CA, respectively). Six kinds of growth medium were applied to grow *E. coli*, resulting in slow to fast cellular growth: M9-glycerol, M9-glucose, M9-glucose-1×CA, M9-glucose-5×CA, LBK, LBKG. SOB medium, consisting of 20 g bacterial tryptone, 5 g yeast extract, 0.584 g NaCl, 0.186 g KCl (J. T. Baker), 2.033 g MgCl_2_·6H_2_O (J. T. Baker), and 2.465 g MgSO_4_·7H_2_O (J. T. Baker) in one liter of deionized water, was used in preparation of electrocompetent cells. SOC medium, SOB medium supplemented with filter-sterilized glucose (180 g/L), was applied to restore cells after electroporation. To achieve a higher transformation efficiency in promoter library construction, we used Recovery Medium (LGC Biosearch Technologies) to revive electrocompetent cells instead of SOC medium. Solid medium was made by supplementing one liter of liquid medium with 20 g agar (BioShop).

### Plasmid and library construction

All primers and oligonucleotides were synthesized by Integrated DNA Technologies and listed in Supplementary Table 1. Most plasmids were constructed by inverse PCR, while some were made by restriction digestion and ligation. Plasmid features and construction procedures are delineated in Supplementary Table 2. Primers for inverse PCR were 5’ phosphorylated using T4 polynucleotide kinase (NEB). Inverse PCR was conducted using Phusion DNA polymerase (Thermo Fisher Scientific). DpnI restriction enzyme (NEB) was employed to remove template plasmids. T4 DNA Ligase (NEB) was utilized to circularize plasmids. After purification by the Zymo DNA Clean & Concentrator kit, the ligation product was transformed into *E. coli* using the BIO-RAD MicroPulser electroporator. Electrocompetent cells were prepared following established protocols^34^. The promoter backbone and the ribosome binding site on the template plasmids for constructing the PL, SP16, SP17, SP18, and DL libraries were adjusted to prevent growth inhibition by excessive GFP expression while providing a wide dynamic range of fluorescence signals for FACS experiments^34^. Primers for library construction were pre-phosphorylated and HPLC-purified. PL, SL16, SL17, SL18, and DL libraries were constructed by inverse PCR of template plasmids, followed by ligation, and transformation into *E. coli* (Supplementary Table 2). Q5 DNA polymerase (NEB) and two inverse primers, each containing randomized nucleotides, were applied to amplify template plasmids. Following DpnI treatment, T4 DNA Ligase was utilized to circularize plasmids. After DNA purification, ligation products were mixed with 180 µL of freshly made electrocompetent cells and divided into six equal aliquots for electroporation. All transformed cells were pooled and revived in 6 mL of Recovery Medium at 37°C and 225 rpm for 1 hour, transferred to 30 mL of LBK medium, and incubated at 37°C and 225 rpm overnight. The transformation efficiency for library construction was ≥ 5×10^7^ colony forming units. Cells in the overnight culture were collected by centrifugation, resuspended in 1 mL of PBS containing 25% glycerol (v/v), aliquoted into 20 portions, and stored at-80°C as the inoculum for sort-seq experiments.

### Measurement of cell growth and GFP expression

Cellular GFP fluorescence was quantified by a BD FACSJazz cell sorter. Each experiment began with inoculating 1 μL of *E. coli* frozen stocks into 2 mL of growth medium incubated at 37°C and 225 rpm overnight. The next day, 5 μL of the pre-culture was transferred to 2 mL of growth medium, incubated and monitored in a Tecan Spark 10M microplate reader. When OD_600_ reached 0.55-0.65, 100 μL of culture was mixed thoroughly with 900 μL of pre-chilled phosphate-buffered saline (PBS) and stored on ice. For each strain, the cell sorter quantified GFP fluorescence (excitation/emission: 488 nm/513 ± 8.5 nm) of 50,000 to 200,000 cells in three replicates.

### Sort-seq experiment

One aliquot of the promoter library frozen stock was revived in 20 mL LBKG medium. Meanwhile, spike-in variants, promoters with known sequences and expression levels ( Supplementary Table 3), were revived in 1 mL LBKG medium. Promoter libraries and spike-in variants were incubated overnight at 37°C and 225 rpm. Subsequently, 50 μL and 2.5 μL of the library and spike-in precultures were transferred to 20 mL and 1 mL LBKG medium, respectively. Both were grown at 37°C and 225 rpm until their OD_600_ reached 0.55-0.65. All spike-in variants were evenly pooled and added to the library in a 1:2500 ratio (v/v). This mixture was diluted ten times with PBS and stored on ice. FACS experiments were performed by a BD FACSJazz cell sorter, and cells were maintained at 4°C throughout. The photomultiplier tube of the cell sorter was calibrated based on spike-in variants pWS19, LP02v14, and LP03v4. The fluorescence distribution of each library was divided into 7-8 ranks, numbered in ascending order according to the GFP fluorescence intensity (Supplementary Table 4). A total of 1.5×10^7^ cells were collected, with the number of cells collected for each rank proportional to its relative abundance in the library. After FACS, cells in each rank were immediately grown in 10 mL LBK medium at 37°C and 225 rpm overnight. Subsequently, the plasmids of each rank were extracted using the Qiagen Plasmid Miniprep Kit. To inspect the FACS purity of each rank, 5 μL of overnight cultures were inoculated into 2 mL LBKG medium incubated at 37°C and 225 rpm.

The fluorescence distribution was examined by the cell sorter when its OD_600_ reached 0.55-0.65 (Fig. 2b and Extended Data Fig. 6a). Promoter sequences in each sorting rank were determined by the Illumina NovaSeq 6000 system. Sequencing library preparation followed the Illumina two-step PCR procedure. A 0.3-kb fragment, including the promoter, 5′ untranslated region, and partial *gfp* gene, was amplified by amplicon PCR using KAPA HiFi DNA polymerase (Roche), along with equal-volume mixed forward primers ILp11, ILp11a, ILp11b, and ILp11c and the reverse primer ILp12. Amplicon PCR products were purified by AMPure XP Beads (Beckman) and then subjected to index PCR using KAPA HiFi DNA polymerase and a unique pair of index primers from the Nextera XT Index Kit (Illumina). Index PCR products were purified by AMPure XP Beads, whose DNA concentrations were measured by a Qubit fluorometer (Thermo Fisher Scientific). Index PCR products were pooled based on the proportion of each rank in the library. The multiplexed sample was subjected to 150-bp paired-end sequencing. We set FastQC quality score Q ≥ 25 to filter out poor sequencing results.

### Using sort-seq data to estimate promoter strength

The occurrence of each promoter variant was counted in each sort-seq rank. Let *F_v_* (*n*) denote the frequency of promoter variant *v* in rank *n*, where Σ*_n_ F_v_* (*n*) = 1. The rank mean *R_v_* of variant *v* was calculated as *R_v_ =* Σ*_n_ n*·*F_v_* (*n*). The rank means of spike-in variants were computed in the same manner. Let *E_v_* denote RFU on the Log_10_ scale and *E_v_* = *a*·*R_v_* + *b*. Parameters *a* and *b* were obtained by performing a linear regression of *R_v_* and *E_v_* of the spike-in variants (Supplementary Table 5). The rank mean of a promoter variant was then converted into its promoter strength using *E_v_*. We set a minimal read count threshold for promoter variants (PL: 20, SL16: 8, SL17: 23, SL18: 31, DL: 9) such that the Pearson’s correlation coefficients of two sort-seq experimental replicates were ≥ 0.9 (Extended Data Fig. 6b,c). To select data with high reproducibility for model training, we further required variants in the PL library to be detected in at least two sort-seq ranks. (Fig. 2b). We combined data from two sets of sort-seq replicates, averaging promoter strength for variants present in both replicates, for subsequent analysis.

### RNA extraction and quantitative PCR

*E. coli* was grown in 10 mL of LBKG medium at 37°C and 225 rpm overnight. When OD_600_ reached 0.55-0.65, cells were harvested by adding 1/10 the volume of a growth-stopping solution (5% Tris-EDTA saturated phenol and 95% ethanol), followed by centrifugation at 9,000 × g for 10 minutes at 4°C. Total RNA was extracted using the QIAGEN RNeasy Mini Kit, followed by removal of genomic DNA with the Turbo DNA-free Kit (Thermo Fisher Scientific). cDNA was synthesized by the High-Capacity cDNA Reverse Transcription Kit (Thermo Fisher Scientific). For each variant, mRNA extraction and cDNA synthesis were performed twice independently. The primer pairs for detecting the *gfp* and *gapA* transcripts were YLEp5/YLEp6 and YLEp7/YLEp8, respectively. Real-Time PCR of each cDNA sample was performed in two replicates with the iQ SYBR Green Supermix (Bio-Rad) on a CFX Connect Real-Time PCR System (Bio-Rad). Only measurements with the standard deviation of cycle threshold values ≤ 0.2 were retained for analysis. The *gapA* housekeeping gene served as the reference for data normalization. mRNA levels were calculated based on an establxished method^34,74^.

## Data availability

The raw sequence reads of the PL, SL16, SL17, SL18, and DL libraries are deposited under National Center for Biotechnology Information BioProject ID PRJNA1049603, PRJNA1049606, and PRJNA1138934.

## Code availability

The code of the promoter prediction model ‘Promoter Scanner’ is available at GitHub (https://github.com/AntonySTKuo/Promoter-scanner/tree/main).

## Acknowledgments

We thank the National Taiwan University College of Life Science Technology Commons for instrument support, Uwe Sauer, Chris Marx, Sheng-Hong Chen, Jun-Yi Leu, and Sen-Lin Tang for insightful discussions. This research is supported by National Taiwan University grants (104R104034, 111L7842, 112L7834, 113L4000-1) and Taiwan National Science and Technology Council grants (110-2311-B-002-008, 111-2311-B-002-020, 112-2311-B-002-011, 113-2311-B-002-008).

## Author contributions

H.H.D.C. conceived the idea, acquired funding, supervised the project, and wrote the manuscript with input from all authors. S.T.K., J.K.C., C.C. and H.H.D.C. developed experimental and analytical methods. S.T.K., J.K.C., C.C., W.Y.S., C.H., S.W.L. and H.H.D.C. led the investigation. H.H.D.C. and S.T.K analyzed and visualized the data.

## Correspondence and requests for materials

should be addressed to Hsin-Hung David Chou.

## Additional files

Supplementary Data 1

Supplementary Information

Extended Data Figs. 1-7

Supplementary Tables 1-5

References for Supplementary Information

